# Nutrient-sensitive reinforcement learning in monkeys

**DOI:** 10.1101/2021.06.20.448600

**Authors:** Fei-Yang Huang, Fabian Grabenhorst

## Abstract

Animals make adaptive food choices to acquire nutrients that are essential for survival. In reinforcement learning (RL), animals choose by assigning values to options and update these values with new experiences. This framework has been instrumental for identifying fundamental learning and decision variables, and their neural substrates. However, canonical RL models do not explain how learning depends on biologically critical intrinsic reward components, such as nutrients, and related homeostatic regulation. Here, we investigated this question in monkeys making choices for nutrient-defined food rewards under varying reward probabilities. We found that the nutrient composition of rewards strongly influenced monkeys’ choices and learning. The animals preferred rewards high in nutrient content and showed individual preferences for specific nutrients (sugar, fat). These nutrient preferences affected how the animals adapted to changing reward probabilities: the monkeys learned faster from preferred nutrient rewards and chose them frequently even when they were associated with lower reward probability. Although more recently experienced rewards generally had a stronger influence on monkeys’ choices, the impact of reward history depended on the rewards’ specific nutrient composition. A nutrient-sensitive RL model captured these processes. It updated the value of individual sugar and fat components of expected rewards from experience and integrated them into scalar values that explained the monkeys’ choices. Our findings indicate that nutrients constitute important reward components that influence subjective valuation, learning and choice. Incorporating nutrient-value functions into RL models may enhance their biological validity and help reveal unrecognized nutrient-specific learning and decision computations.

## INTRODUCTION

According to the influential Reinforcement Learning (RL) framework, animals learn by updating reward values based on experience and chose by comparing these values between options^1^. The RL framework has been critical for identifying fundamental learning and decision variables that guide animals’ behaviour, including object values and action values, which provide essential decision inputs, and the reward prediction error, which updates values from experience. Direct physical implementations of these theoretical constructs have been discovered in the activity of neurons in primate dopamine neurons^2–5^, striatum^6,7^, amygdala^8,9^, and prefrontal cortex^10–13^. Despite its broad explanatory power, the RL framework does not explain how learning and choice depend on specific reward properties. For example, nutrients are biologically critical, intrinsic components of food rewards, and an animal’s survival depends on its ability to make adaptive food choices that acquire specific nutrients. Investigating how nutrient rewards influence learning and choice could not only enhance the biological validity of RL models. It may also guide the discovery of so-far unrecognized nutrient-specific learning and decision computations, and their neuronal implementations.

Because nutrients are mainly acquired from food intake, an animal’s ability to adapt its food choice to changing nutrient availabilities critically determines its nutrient balance and long-term health. To optimize nutrient intake, foraging animals adapt their feeding patterns in response to regional and seasonal variations of food resources^14–16^. For instance, monkeys spend more time in food patches associated with a high probability of nutritious foods (e.g., nuts) while ignoring more frequent low-nutrient foods (e.g., leaves). Primates, including humans, also exhibit individual subjective preferences for specific nutrients and sensory food qualities to regulate nutrient intake^17–24^. Thus, ecological data suggest that animals consider both the nutritional value of food and the food’s availability. However, the specific learning and decision computations underlying such nutrient-sensitive food choices remain unclear. Here, we examined the food choices of rhesus monkeys (*Macaca mulatta*) in a dynamic foraging task that involved choices between rewards with different nutrient (fat, sugar) components under varying reward probabilities.

Previous studies examined how monkeys adapt to changing reward probabilities^9–13,25–27^. In probabilistic learning tasks, monkeys track the high-probability option based on past choices and reward outcomes and distribute their choices according to the reward probability of both options. This learning strategy has been modelled by linking subjectively weighted recent rewards to current choices (‘reward history’) using logistic regression^25,26^ and by dynamic updating of option values based on reward outcomes via RL mechanisms^1^. We followed these approaches and examined whether monkeys assigned higher value to more nutritious foods during learning and learned faster from high-nutrient rewards.

First, we characterized monkeys’ nutrient preferences and learning during probabilistic reward-based choices. If the monkeys preferred specific nutrients, they should choose high-nutrient rewards more frequently and track their changing probability more closely to maximize intake of the specific nutrient. We recently showed in a nutrient-choice task without learning requirement that macaques’ choices reflect underlying, stable nutrient-value functions^22^. Accordingly, we hypothesized that nutrient-value functions also govern choices during probabilistic reward learning.

Next, we examined whether monkeys demonstrated nutrient-specific learning. We followed established approaches for characterizing the integration of past reward experiences into subjective values using logistic-regression and RL frameworks^10,11,25,26^ to examine whether nutrient preferences modulated reward learning. To account for nutrient-specific learning, influences of recent reward and choice histories on current choice should be higher for high-nutrient reward. Accordingly, the value function in a formal RL model should incorporate higher preferences for high-nutrient rewards (‘nutrient-value function’). In addition, the animals may assign higher weights to reward outcomes with particular nutrient content, as reflected by influences on learning rate (‘nutrient-specific learning rates’).

Finally, based on behavioral evidence for nutrient-sensitive reinforcement learning, we propose candidate neuronal mechanisms necessary to implement nutrient-specific learning and decision computations, as a framework to guide future neurophysiological recordings.

## RESULTS

Two monkeys performed in a dynamic foraging task to obtain different nutrient-defined liquid rewards (**Fig. 1A**). In each choice trial, the monkeys were presented with two visual cues from a set of four, chose between the two cues, and received either a large amount (‘rewarded’) or a small amount (‘non-rewarded’) of the cue-associated liquid reward, depending on a prespecified reward probability (*p*). We used new, untrained visual cues in each session to avoid influences of prior experience. Session-specific visual cues were each associated with one of four different rewards; cue-reward associations were fixed within each session. To examine whether fat and sugar biased learning from reward outcomes, we used liquid rewards from a 2 × 2 factorial design with fat and sugar levels as factors (**Fig. 1B**; LFLS: low-fat low-sugar; HFLS: high-fat low-sugar; LFHS: low-fat high-sugar; HFHS: high-fat high-sugar). At the start of each session, two rewards (LFLS/HFHS or LFHS/HFLS) were associated with a high probability of obtaining a large reward (*p* = 0.8), and the other two rewards were associated with a low reward probability (*p* = 0.2) (**Fig. 1C**, block A or block B). We reversed the reward probabilities every 100 trials throughout the session (*p* = 0.2 **→** 0.8; *p* = 0.8 **→** 0.2) to encourage continual learning from reward outcomes (**Fig. 1D**). Notably, this design offered the monkeys equal availability of fat and sugar in all choice trials irrespective of block type because there were always two high-probability and two low-probability options for both high-fat and high-sugar rewards. All liquids were matched in flavour (blackcurrant or peach) and other ingredients (protein, salt, etc); therefore, differential learning and choice patterns could be attributed to the nutrient content of the rewards.

**Fig. 1.**
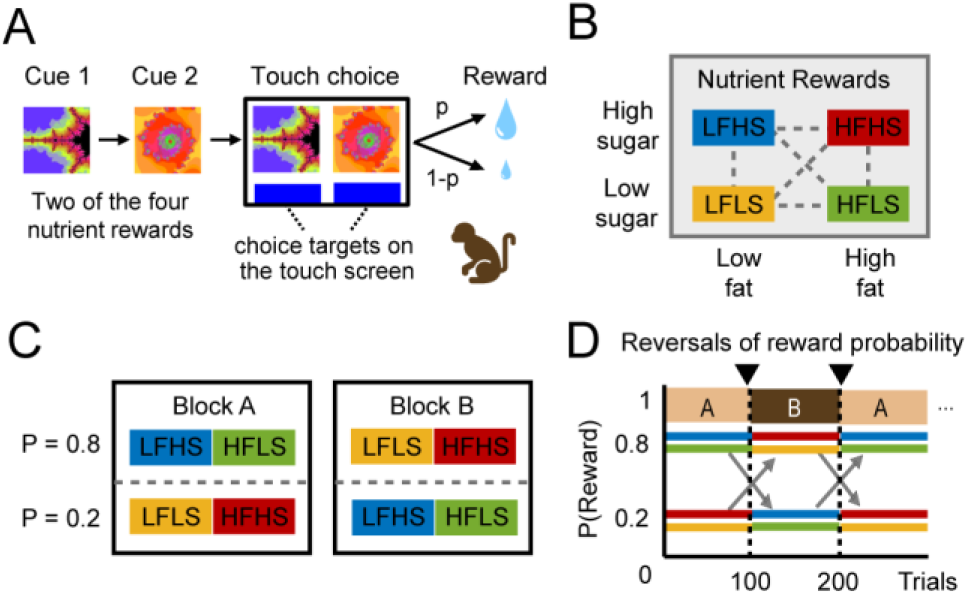
Nutrient foraging task. A) Task structure. In each trial, two visual cues appeared sequentially on a touch screen before reappearing in a left-right arrangement as choice targets. Following the touch choices, the monkeys received the liquid reward associated with the chosen cue. The amount of the delivered reward depended on a prespecified reward probability (p). B) Four types of liquids with 2 × 2 factorial fat and sugar levels were offered to the monkeys: the low-fat low-sugar (LFLS) liquid, the high-fat low-sugar (HFLS) liquid, the low-fat high-sugar (LFHS) liquid, and the high-fat high-sugar (HFHS) liquid. C) Reward probabilities associated with the different reward types reversed between blocks of trials in a testing session. In block A, LFHS and HFLS were associated with a high probability (*p* = 0.8) of receiving the large reward, LFLS and HFHS were associated with a low probability (*p* = 0.2) of large reward; these probabilities reversed in block B. D) Each session started with either block A or block B and the reward probabilities reversed every 100 trials between the two block types, with typically 3-5 reversals per session.

### Nutrients bias reward learning and food choices

The behaviour in two example sessions (**Fig. 2A**) showed that both monkeys exhibited preferences for specific nutrients while tracking changing reward probabilities. Monkey Ya’s choices (**Fig. 2A**, top) were dominated by a general preference for high-sugar rewards, with a smaller impact of reward probability on choice. Specifically, monkey Ya chose the high-sugar rewards frequently even when they were associated with a lower probability of obtaining a large reward amount; in addition, choice frequencies tracked changing reward probabilities, particularly for the high-sugar rewards. By contrast, monkey Ym’s choices (**Fig. 2A**, bottom) reflected both a preference for high-nutrient content and a strong dependence on reward probability. Specifically, within a given trial block, monkey Ym preferred high-nutrient rewards over low-nutrient rewards with matched reward probabilities (compare red and yellow curves) but would reduce his choices for the high-nutrient reward when it was associated with a relatively lower reward probability.

**Fig. 2.**
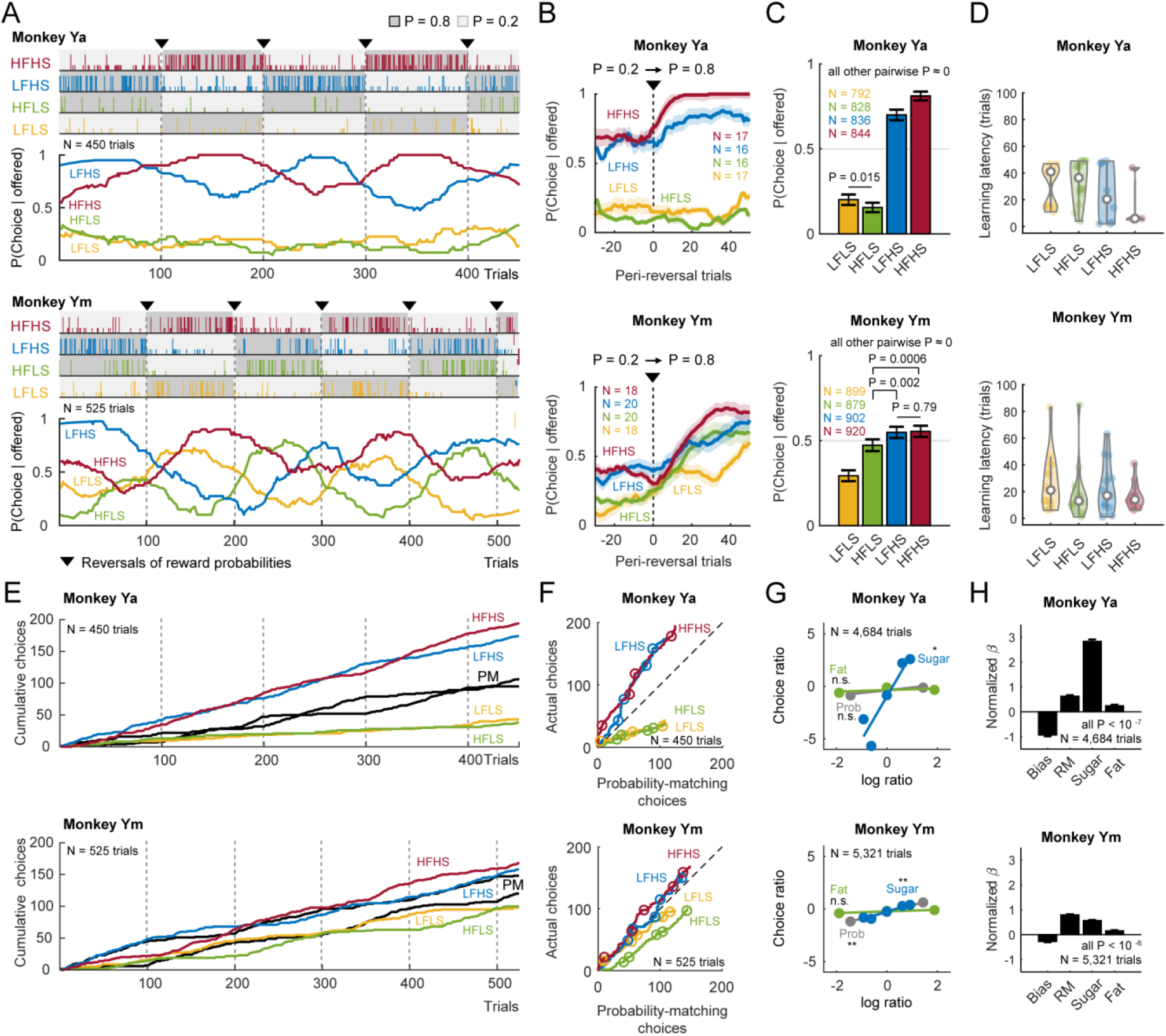
Nutrient-sensitive learning and choice in monkeys. A) Choices and reward outcomes in a single session for monkey Ya (top) and monkey Ym (bottom). Each tick mark represents a choice of a specific reward type; long marks indicate large reward outcome, short marks indicate small reward outcomes. Reward types in dark-gray blocks were associated with high reward probability (*p* = 0.8) and those in light-gray blocks were associated with low reward probability (*p* = 0.2). Choice curves showed running-average choice patterns of each reward. B) Learning curves. Mean running-averaged choice frequencies aligned to probability reversals (*p* = 0.2 **→** 0.8) indicate how choices depend on both reward-probability changes and nutrient content. (N: number of tested sessions). C) Reward preferences. Average choice frequencies indicate preferences among the four reward types. The choice frequencies were computed after sessions were truncated, including only probability-balanced trials for all reward types. (mean ± s.e.m.) (N: number of trials). D) Learning latency. The number of trials from probability reversal to the first significant change point in the cumulative choice record (see *Methods*) indicates latency to adapt choices after probability changes. E) Monkeys’ single-session cumulative choice records deviate from the pure probability-matching strategy. PM: probability-matching choice strategy, calculated by matching choices to the past ratio of large/small rewards, irrespective of reward type. F) Direct comparisons of monkeys’ choices with probability-matching choices. Circles indicate probability reversals. G) Nutrient-sensitive matching behavior. Correlations of choice ratios with fat, sugar, and probability ratios, respectively. H) Normalized regression coefficients of probability ratios, fat ratios, and sugar ratios on choice ratios. (mean ± s.e.m.).

The patterns observed in single sessions were also observed in averaged data across sessions. Overall, the monkeys’ choice probabilities increased when reward probabilities switched from low (*p* = 0.2) to high (*p* = 0.8), as evident by averaged choice probabilities around probability-reversal points (**Fig. 2B**). Importantly, the monkeys responded differently to probability changes for rewards that differed in fat and sugar content, with more pronounced probability increases for high-nutrient rewards and specifically high-sugar rewards (**Fig. 2B**). When reward probabilities were stable (between reversal points), monkey Ya showed a strong preference for the high-sugar rewards irrespective of fat level, whereas monkey Ym showed graded preferences for both high-fat and high-sugar rewards over the low-nutrient option (**Fig. 2C**). Immediately following the probability reversals, the monkeys had shorter learning latencies for high-nutrient rewards: they adjust their choices more quickly to the changed reward probabilities when high-sugar and high-fat rewards were offered, which indicated that learning was sensitive to the nutrient content of reward outcomes (**Fig. 2D**). Thus, the monkeys preferred high-nutrient rewards, tracked changing reward probabilities in a nutrient-dependent manner, and learned faster from high-nutrient reward outcomes.

The preferences for fat and sugar biased the monkeys’ choices away from a pure probability-matching (PM) strategy, which predicted distributed choices according to the relative frequency of receiving large rewards from each option. In the two example sessions, choices for the high-sugar rewards accumulated more rapidly than predicted by the PM strategy, whereas choices for low-sugar rewards accumulated more slowly (**Fig. 2E**). Specifically, compared to the PM strategy, monkey Ya significantly over-matched the high-sugar rewards and under-matched the low-sugar rewards, irrespective of reward-fat level. These patterns were much less pronounced in monkey Ym (**Fig. 2F**). Specifically, the choice ratios of monkey Ya were dominated by the sugar ratios but those in monkey Ym were jointly determined by the probability ratios and sugar ratios (**Fig.2G**). Multiple regression confirmed that, in addition to the probability ratios, both the fat and sugar ratios significantly influenced the choice ratios (**Fig. 2H**). Notably, both monkeys’ choices were explained by similar effect sizes of the probability ratios and the fat ratios. However, the effects of sugar ratios were particularly strong in monkey Ya but slightly weaker than the influences of probability ratios in monkey Ym.

Taken together, these results suggested that the specific nutrient composition of food rewards and the animals’ individual preferences for sugar and fat biased learning and choice.

### Nutrient-specific reward history and choice history influence monkeys’ choices

One strategy to respond to unsignaled changes in reward probabilities is to choose based on recent choices and reward outcomes. Because the choice outcomes reflect the underlying reward probability, this strategy adapts choices to the changing reward probabilities and can help to optimize reward rate and nutrient-intake levels. Consistent with these notions, we found that monkey Ym tended to repeat his choices, particularly after receiving a large reward on the previous choice; this effect was evident across all reward types (**Fig. 3A,** right). By contrast, the tendency to repeat choices was less pronounced for the low-sugar rewards in monkey Ya (**Fig. 3A**, left). This result suggested that both recent choices and the reward outcomes increased choice repetition, but the influences depended on individual nutrient preferences.

**Fig. 3.**
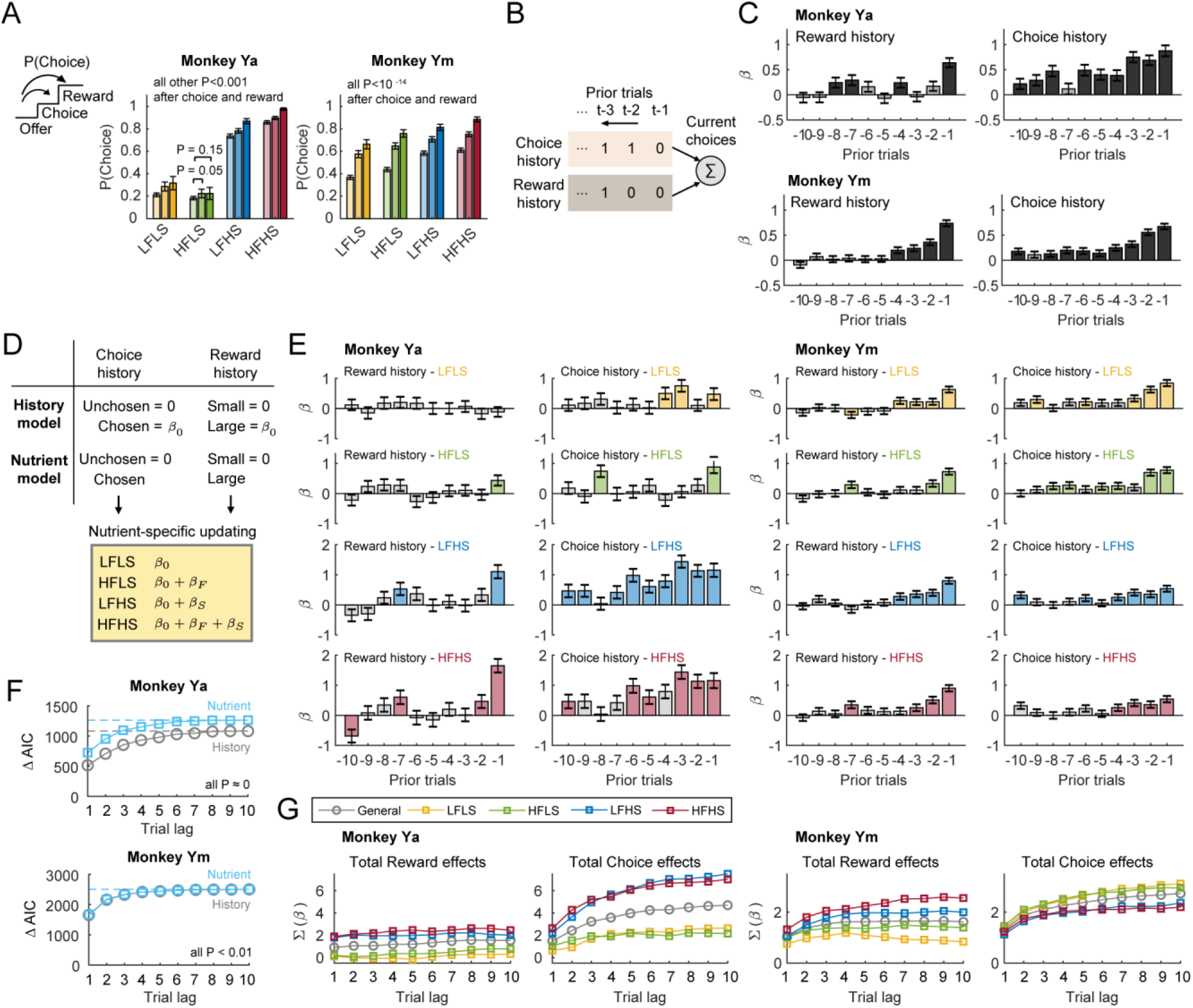
Nutrient-specific reward and choice histories influence monkeys’ choices. A) Choice probabilities for different rewards depended on recent experience. Choice probabilities when the same reward was chosen on the previous trials (‘choice’), when a large reward was received on the previous trial (‘reward’), and irrespective of last-trial choice and reward outcomes (‘offer’). B) History model for explaining the choice. The history model included regressors for choice history and reward history, with choice history = 1 (chosen) or 0 (not chosen) and reward history = 1 (rewarded) or 0 (non-rewarded) for the past 10 trials. C) Results, history model. Logistic-regression coefficients show influences of reward (left) and choice history (right) on current choices. (mean±s.e.m.; dark-gray bars: P < 0.05; light-gray bars: non-significant) D) Nutrient model for explaining choices. In contrast to the assumption of uniform history effects across reward types in the history model, the nutrient model examined nutrient-specific history effects by including additional nutrient-specific history regressors. E) Results, nutrient model. Nutrient-specific logistic regression coefficients for current choices. (P < 0.05, yellow: LFLS; green; HFLS; blue: LFHS; red: HFHS; light-gray bars: non-significant) F) Model performances and history lengths. Model performance improved with history length based on ΔAIC = AIC (trial lag = 0) – AIC (trial lag = *i*, *i* = 1,2,…,10). AIC = Akaike Information Criteria. History length-matched nutrient models and history models were compared using the loglikelihood test. Higher ΔAIC values indicated that the nutrient model outperformed the history model in all history length-matched comparisons. G) Aggregated effects of reward and choice history increased with history lengths and reflected nutrient composition, indicated by the cumulative reward or choice history regression coefficients over recent trials.

To formally characterize the learning from recent choices and reward outcomes, we modelled the trial-by-trial choices in a logistic regression model (history model, see *Method*) that accounted for whether the option was chosen in previous offers (choice history) and whether the previous choices were rewarded (reward history) (**Fig. 3B**). The regression coefficients showed that both the choice and reward history reinforced current choices and that these effects decayed for more remote past trials (**Fig. 3C**). Given the monkeys’ preferences for fat and sugar, we next examined whether these reward- and choice-history effects also depended on the nutrient composition of reward outcomes and choice offers. We tested this possibility by including nutrient-history interaction regressors in the history model (nutrient model, see *Method*). These interaction terms would capture any additional reinforcing effects from specific nutrients by decomposing the aggregated reward and history effects into the effects of baseline low-nutrient liquid (*β*_0_), high-fat content (*β_F_*), and high-sugar content (*β_S_*), depending on the fat and sugar levels of the offered reward types (**Fig. 3D-E**). Larger history regression coefficients for sugar compared to fat suggested that recently obtained high-sugar reward outcomes had a stronger impact on current-trial choice than recently obtained high-fat rewards in both monkeys. However, the two monkeys differed in their tendency to repeat choices for high-fat and high-sugar liquids, as indicated by the nutrient-specific choice-history coefficients. Monkey Ya repeated the high-sugar choices more frequently than choices for low-nutrient rewards and high-fat rewards. By contrast, monkey Ym repeated choices slightly less frequently for the high-sugar rewards. Importantly, although the explanatory power of both models increased with history length, the nutrient model outperformed the history model in all history length-matched comparisons (**Fig. 3F**). These history effects showed distinct temporal dynamics in the two monkeys although they both decayed either in the history model or the nutrient model (**Fig. 3G**).

These results indicated that both monkeys’ choices depended on the recent histories of obtaining and choosing rewards with specific nutrient content.

### Reinforcement learning based on nutrient-specific values

The temporal dynamics of the nutrient-specific reward- and choice-history effects suggested that the monkeys constantly updated their choices based on recent choices and reward outcomes. RL models that update trial-by-trial reward values for each option based on the reward outcomes are well-suited to model such adaptive choices. However, canonical RL models typically do not account for the nutrient composition of food rewards, and accordingly cannot explain the presently observed nutrient preferences and nutrient-specific learning effects. Therefore, we developed a nutrient-sensitive RL model that incorporated subjective nutrient values to model how specific nutrients (fat, sugar) differentially influenced the trial-by-trial updating of expected reward values and their influence on the choice (**Fig 4A**). Instead of updating the value of the chosen reward with a binary reward outcome, our model updated reward values based on the nutrient composition of each reward type as given below,

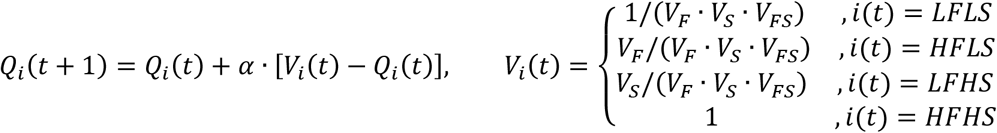

where the value for reward *i, Q_i_*, was updated depending on the chosen reward type on trial *t, i*(*t*) and its nutrient-specific reward value, *V_i_*(*t*). *V_F_, V_S_*, and *V_FS_* denoted the subjective value of high-fat content, high-sugar content, and their interaction, respectively, on the common scale of the low-nutrient reward value. Therefore, any nutrient value larger than 1 suggested a preference for the specific nutrient; values for larger than 1 indicated supra-additive values of fat and sugar. Without loss of generality, we normalized all reward values to the highest nutrient value, (*V_F_* · *V_S_* · *V_FS_*), to constrain all reward values between 0 and 1. For the unchosen and unoffered rewards, we allowed the values to decay as follows,

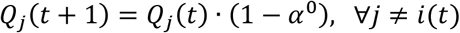

where the values of the unchosen and unoffered rewards, *Q_j_*(*t*), were discounted according to a forgetting rate (*α*^0^), which would be 0 for perfect (but biologically implausible) value memory.

**Fig. 4.**
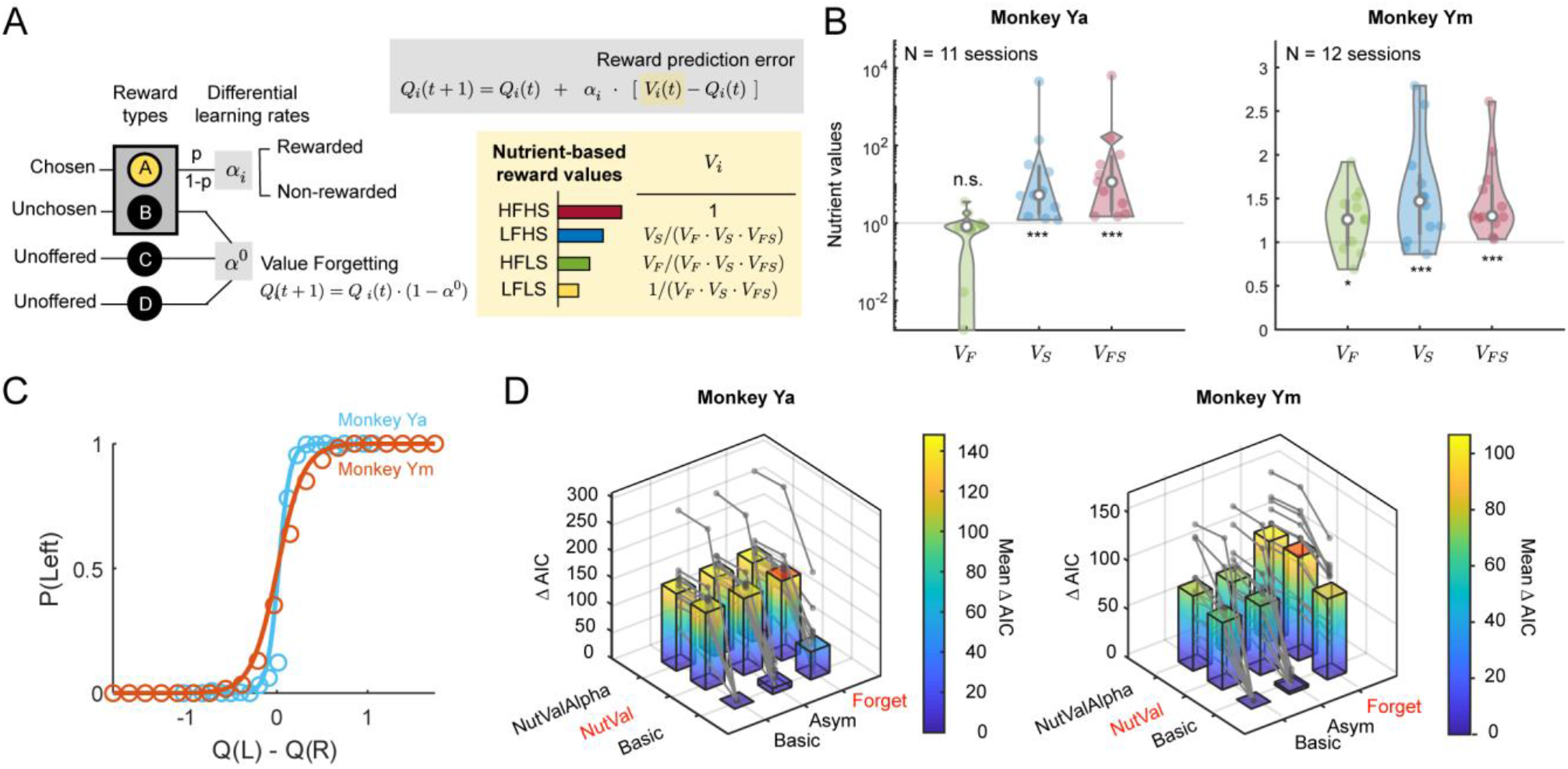
Nutrient-sensitive reinforcement learning models. A) The nutrient-value RL model (NV-RL model). Reward values were updated based on the nutrient-specific values, *V_i_*(*t*), with *V_F_* indicating values for the high-fat content, *V_S_* for the high-sugar content, *V_FS_* for the high fat-sugar combination, and 1 for the low-nutrient value reference. We normalized all reward values to the highest reward value (*V_F_* · *V_S_* · *V_FS_*) to constrain all reward values between 0 and 1. B) Nutrient-specific reward values. The distributions of fitted nutrient-specific reward values across trials (monkey Ya: log scale; monkey Ym: linear scale). All reward values were tested against equal values for all reward types (nutrient values =1), Wilcoxon signed-rank test. C) Nutrient-value functions. Psychometric curves based on integrated values, calculated with the nutrient-value RL model, indicate that both monkeys’ choices depended on nutrient-dependent value differences between choice options. D) Model comparisons. The main nutrient-value RL model (NutVal-Forget model) was systematically compared with alternative RL models involving combinations of differential learning rates (*NutVal* = nutrient-specific values; *NutValAlpha* = nutrient-specific values + learning rates, Figure S2A) and nutrient-specific parameters (*Asym* = independent learning rate for the non-rewarded chosen option; *Forget* = value-forgetting for unchosen and unoffered options). Models were compared using Akaike Information Criterion (AIC). All model AICs were subtracted from the AIC of the basic RL model (Δ*AIC* = *AIC_basic_* – *AIC*) for comparison. The higher mean Δ*AIC* indicated better model performance (red: the best fitting model).

The results of fitting this nutrient-sensitive RL model to each monkeys’ choices and reward outcomes in each session confirmed that both monkeys assigned higher values to the high-sugar choice options and that monkey Ym assigned higher value to fat but monkey Ya did not (**Fig. 4B**). The high-fat high-sugar reward was also valued higher than the low-nutrient reference, but the fat values and the sugar values did not show supra-additive effects in monkey Ya but negative interactions in monkey Ym when determining the reward values (**Fig. S1**). The model-derived subjective values for fat and sugar accurately predicted the monkeys’ choices (**Fig. 4C**). The nutrient-sensitive RL model outperformed alternative RL models involving combinatorial differential learning rates and nutrient-specific parameters (**Fig. 4D;** see *Methods*). Notably, there was no evidence for nutrient-specific learning rates but only a significant but small forgetting rate for monkey Ym (**Fig. S2**).

Thus, the monkeys’ stochastic choices for rewards with specific nutrient compositions were well explained by a nutrient-sensitive RL model that assigned nutrient-specific values to reward outcomes.

### Value updating based on distinct sugar and fat value components

The nutrient-sensitive RL model implied that the animals can independently track values for specific fat and sugar nutrients, and integrate them into a scalar value that guided choices. To better understand the dynamics of this nutrient-specific value tracking and updating, we modelled the dynamic learning of individual nutrient values in a nutrient prediction error-based RL model (NPE-RL) in which the reward value on trial *t, Q_i_*(*t*), was jointly determined by individual fat value and sugar value components (**Fig. 5A,** see *Methods*).

**Fig. 5.**
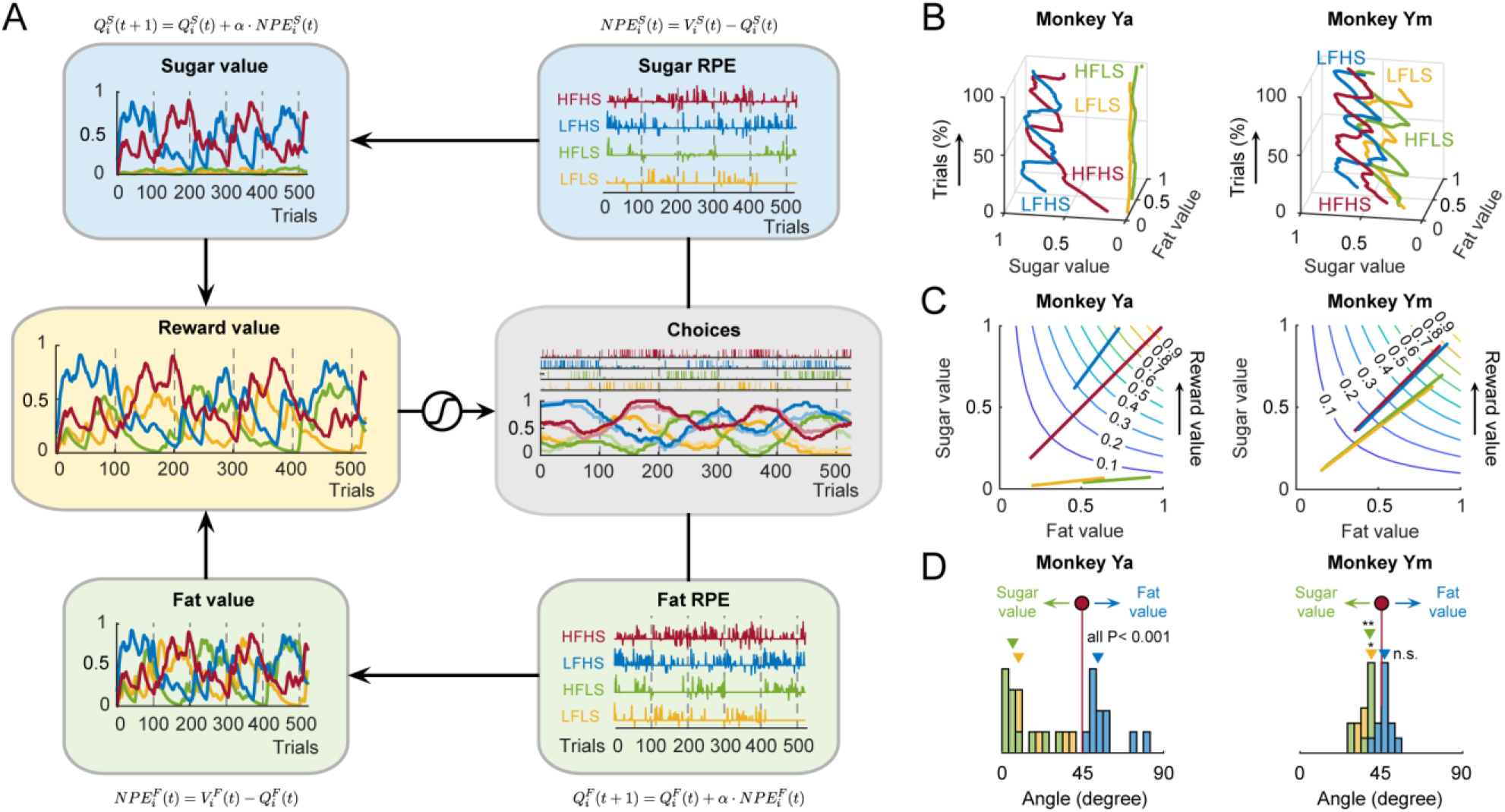
Dynamics of sugar and fat value components in nutrient-sensitive reinforcement learning. A) Nutrient-specific value updating. Nutrient-specific values for sugar (top) and fat (bottom) were updated based on discrepancies between previous choice outcomes and predicted nutrient rewards (nutrient prediction errors, NPE); sugar and fat values were integrated into composite reward values that guided choices. B) Trajectories of nutrient-specific values within sessions. The value trajectories tracked the evolving reward values and their nutrient components with choice trials. C) Projected reward-value trajectories and iso-value contour curves. Each segment showed the ranges and orientations of the fluctuating reward values in the nutrient value space. The diagonal line represented equal contributions of the nutrient components to the reward values. D) Nutrient sensitivities of reward values. Distributions of the rotating angles quantified the relative changes of nutrient values during reward value updating (Δ*V_S_*/Δ*V_F_*) across sessions. Wilcoxon signed-rank test.

The NPE-RL model characterized how fat and sugar values could (i) separately adapt to changes in reward probabilities as indicated by experienced outcomes, and (ii) flexibly determine the integrated reward values for specific choice options, based on their nutrient composition. Specifically, the fat and sugar RPEs for each reward updated the fat and sugar values, respectively, which were then combined into integrated reward values to guide choices (**Fig. 5A**). Decomposing the reward values into two independent nutrient components revealed each animal’s idiosyncratic sensitivity of reward values to individual nutrient constituents. To illustrate the dynamic, nutrient-specific value updating, we plotted the evolving value trajectories within a session in a space defined by the separate fat and sugar value components (**Fig. 5B**). These trajectories indicated that the updating of reward values in monkey Ya was primarily based on the sugar value component over the fat value component, whereas both fat and sugar value components contributed to value learning in monkey Ym (**Fig. 5B**).

The distinct sensitivities of reward values to specific nutrient components were illustrated by projections of dynamic reward value trajectories onto the nutrient value space, where ‘iso-value contours’ visualized levels of equal reward values (**Fig. 5C**). If reward values were equally sensitive to both the sugar and fat nutrient components, the value trajectories should fall onto the 45-degree diagonal line in nutrient value space. Because we normalized the nutrient values to the HFHS reward, the value trajectory for HFHS would be at the diagonal for both monkeys (**Fig. 5C,** red). However, higher sensitivity to the sugar value components compressed the low-sugar value trajectories along the sugar value axis and rotated the trajectories towards the fat value axis (clockwise); similarly, higher sensitivity to the fat value components rotated the low-fat trajectories towards the sugar value axis (counterclockwise). For example, monkey Ya showed a slight counterclockwise-rotated LFHS trajectory and marked clockwise-rotated low-sugar trajectories, indicating his weak preference for fat and the strong preference for sugar, respectively. In contrast, monkey Ym showed only mild clockwise-rotated low-sugar trajectories and negligible rotation for the LFHS value trajectory, reflecting his mild sugar preference and non-significant fat preference.

The rotating angles of the value trajectories in the nutrient value space quantified the relative changes of sugar and fat values on the value trajectories (Δ*V_S_*/Δ*V_F_*), therefore highlighting the contributions of each nutrient value to the overall reward values. Compared to the steepest 45-degree value gradient, value trajectories with rotating angles larger or smaller than 45 degrees updated reward values with distinct contributions of each nutrient value. Specifically, reward values were mostly updated from the fat values when the angles were smaller than 45 degrees, but more from the sugar values if the angles were larger than 45 degrees. Across sessions, the nutrient-specific contributions of reward values, indicated by the orientations of the value trajectories, recapitulated the subjective nutrient values estimated by the nutrient-sensitive RL model (**Fig. 5D**).

Thus, subjective nutrient-value functions guided the dynamic updating and the integration of reward values based on individual nutrient-specific components.

## DISCUSSION

We investigated monkeys’ choices for different nutrient-defined rewards under varying reward probabilities. We found that the nutrient composition of rewards strongly influenced choices and learning. The animals generally preferred rewards that were high in nutrient content but also showed individual preferences for sugar and fat, consistent with the assignment of subjective values to choice options. The animals’ nutrient preferences affected how they adapted their choices to changing reward probabilities. Specifically, the monkeys learned faster from preferred nutrient-rewards and chose them frequently even under low reward probabilities (i.e., low probability of obtaining large reward amounts). Influences of past rewards on current choice were well described by a reward-history analysis. As in previous studies^11,25^, more recent rewards had a stronger influence on the monkeys’ choices. Critically, we also found that the impact of reward history depended on the nutrient composition of past rewards: the effect of past rewards high in preferred sugar content was stronger compared to that of less preferred low-nutrient or fat rewards. The history of past choices, irrespective of reward outcomes, also had a significant and nutrient-dependent effect on choice, with stronger effects of past choices for preferred nutrient rewards. We proposed a nutrient-sensitive RL model that captured the influences of preferred nutrients on learning and choice. The model updated the value of individual sugar and fat components of expected rewards trial by trial, based on recently experienced rewards, and integrated these components into scalar values that explained the monkeys’ choices. These results suggest that nutrients constitute important reward components that influence subjective valuation, learning and choice, and that canonical RL models can be usefully extended to capture such nutrient-specific values.

Previous studies of reinforcement learning in macaques revealed important influences on learning and choice, including effects of reward and choice history^5,7,11–13,25–27^, the variance of recent rewards^10^, novelty and reward rarity^9,28^, and social observations^8^. Importantly, these studies did not vary the composition of reward outcomes and thus could not test whether specific reward components differentially affected learning and choice. We reasoned that nutrients are biologically critical reward components that are essential for survival and that monkeys should prefer high-nutrient rewards and adapt their choices to optimise nutrient intake. By manipulating the sugar and fat content of our liquid rewards, we confirmed that the monkeys’ learned differently from these different rewards.

Previous studies demonstrated that macaques have sophisticated preferences for different reward types that comply with principles of economic choice theory^2,29–31^ but did not examine how different rewards affect learning. Here we showed that subjective preferences for specific nutrients influenced how monkeys tracked the changing reward probabilities of choice options. Specifically, both animals learned faster from preferred nutrient rewards. Moreover, they based their choices on both subjective valuations of offered reward types and estimates of current reward probabilities. This latter finding confirms the result from a previous study that macaques integrate reward type and probability information to express subjective preferences^29^; different from that study, our monkeys were required to derive probability information from past reward experiences rather than from explicit visual cues.

Crucially, by varying the nutrient composition of rewards, we investigated reinforcement learning and choice for biologically important, universal reward components. Nutrients are basic building blocks of foods that are sensed by dedicated taste and oral-texture mechanisms^22,32–34^ and engage physiological and homeostatic processes^35,36^. Moreover, evidence from ecology and human metabolic sciences points to specific behavioral mechanisms that regulate nutrient intake. For example, ecological studies identify a ‘nutrient-balancing mechanism’ in wild macaques that promotes reproductive and survival success^14,37–39^. In humans, reduced protein in ultra-processed foods increases energy intake by ‘protein leveraging’, a mechanism that regulates food choice to counter protein deficits^35,40–42^. A ‘fat-appetite mechanism’ emerges in human monogenic obesity affecting melanocortin-signalling^21^. We recently showed that in macaques, nutrients and sensory food qualities (taste, viscosity, oral friction) shape human-like economic preferences^22^. Our approach makes a first step towards integrating the influential RL framework with these nutrient-dependent behavioral processes and thus enhance its biological validity.

The concept of nutrient homeostasis in metabolic sciences suggests that internal states modulate nutrient values to guide state-dependent food choices. Recent homeostatic RL models explain the value of rewards as discrepancies between the current state and physiological setpoints^43^. This approach views reward values as physiological signals that serve to maintain homeostasis. However, fat and sugar can be preferred even without corresponding nutrient deficits^22,23,44^; therefore, the hedonic values of foods cannot be explained solely by homeostatic regulations of nutrient deficits. Future experiments could challenge the nutrient states of animals during food choices to estimate empirical nutrient-value functions from state-dependent choice patterns to refine these models.

We described a nutrient-specific learning mechanism that updates value estimates for separate fat and sugar reward-components and integrates this information to guide adaptive food choices. This mechanism implies parallel nutrient valuation systems that detect and evaluate the nutrient components depending on internal states. The neuronal implementation of this mechanism would require neurons that encode individual nutrient values (nutrient-value neurons) and dynamically update these nutrient values via nutrient prediction error signals (**Fig. 6A**). At a neural-network level (**Fig. 6B**), these nutrient-value neurons would extract nutrient-specific features from a food’s sensory properties to guide food choices. Importantly, physiological-state signals could modulate the neural representations of nutrient values to allow for state-dependent valuation of food rewards. Therefore, we propose nutrient-value neurons and nutrient prediction error signals as potential substrates for nutrient-sensitive learning and choice.

**Fig. 6.**
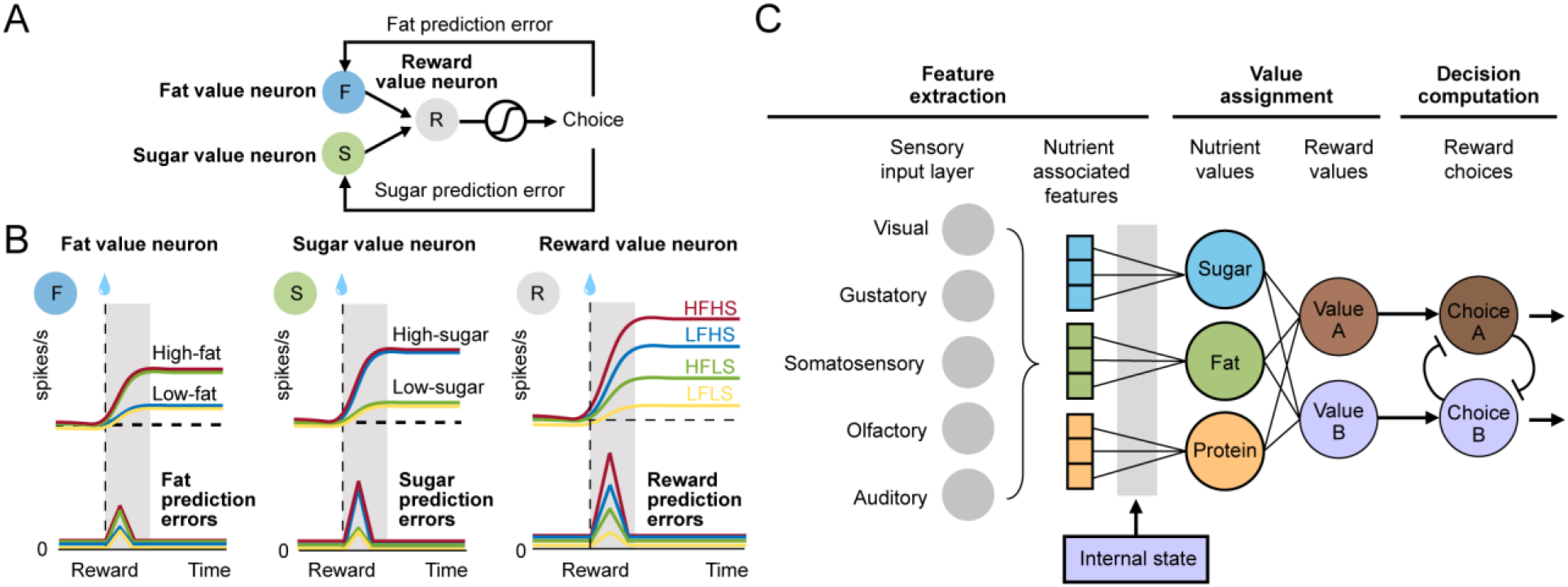
Neuronal mechanisms for nutrient-sensitive reinforcement learning and choice. A) Nutrient-sensitive reinforcement learning architecture. Fat-value neurons (F) and sugar-value neurons (S) each update the fat and sugar components of the value predictions and provide input to the reward-value neurons (R) that code integrated values for decision computations. B) Predicted neuronal responses of fat-value, sugar-value, and reward-value neurons. Fat-value neurons (F) are updated based on fat-specific reinforcement learning (fat prediction errors) from delivered rewards, with higher responses to high-fat compared to low-fat rewards. An equivalent process operates for sugar-value neurons (S). Together, these nutrient-value neurons converge onto reward-value neurons to code scalar value signals in a common currency for downstream decision computations. C) Nutrient-sensitive decision-making neural network. Sensory properties of foods are detected via multiple sensory channels and integrated into nutrient-associated feature representations that determine nutrient values depending on the internal physiological state. Nutrient values then flexibly inform reward values for decision computations, based on the nutrient composition of food rewards.

Our findings within a nutrient-based RL paradigm and our proposed computational framework have implications for value-based learning and decision theories and underlying neural mechanisms. Because nutrients provide energy and serve physiological functions for survival, animal reward systems should be shaped by nutrient availability in the environment and evolved dedicated mechanisms for adaptive nutrient-sensitive decision-making. By decomposing the trial-by-trial reward values that guide reinforcement learning into nutrient-value components, we identified candidate signals that could be encoded by neurons in the reward and decision systems of the primate brain. The midbrain dopamine neurons, orbitofrontal cortex and amygdala participate in decision-making, reinforcement learning, and food evaluation^2,3,8,31,45–48^ and thus constitute suitable targets for testing these hypotheses experimentally.

## METHODS

### Animals

Two adult male rhesus macaques (*Macaca mulatta*) were trained in the study: monkey Ya (weight during the experiments: 17-19 kg, age: 6 years) and monkey Ym (12-13 kg, age 6 years). The animals were trained and tested approximately one to two hours per day and five days per week for 6 months. Both monkeys participated in another nutrient choice study using the same dairy-based nutrient rewards as in this study. The animals were on a standard diet for laboratory macaques, composed of high-protein dry pellets (% calories provided by protein: 30.36%, fat: 13.29%, carbohydrates: 56.34%), dried fruits, seeds, nuts, and fresh fruits and vegetables. We monitored the monkeys’ health condition and body weights to ensure their welfare after introducing high-calorie rewards. No effects of these rewards on the animals’ health were observed. Each testing day, the animals had free access to the standard diet before and after the experiments and received their main liquid intake in the laboratory. The animals’ body weights increased as expected for growing animals.

All animal procedures conformed to US National Institutes of Health Guidelines. The experiments have been regulated, ethically reviewed and supervised by the following UK and University of Cambridge (UCam) institutions and individuals: UK Home Office, implementing the Animals (Scientific Procedures) Act 1986, Amendment Regulations 2012, and represented by the local UK Home Office Inspector; UK Animals in Science Committee; UCam Animal Welfare and Ethical Review Body (AWERB); UK National Centre for Replacement, Refinement and Reduction of Animal Experiments (NC3Rs); UCam Biomedical Service (UBS) Certificate Holder; UCam Welfare Officer; UCam Governance and Strategy Committee; UCam Named Veterinary Surgeon (NVS); UCam Named Animal Care and Welfare Officer (NACWO).

### Experimental Design

#### Nutrient rewards

We prepared nutrient-controlled liquids with 2 × 2 fat and sugar levels to examine whether fat and sugar biased learning from reward outcomes (**Fig. 1B**; LFLS: low-fat low-sugar; HFLS: high-fat low-sugar; LFHS: low-fat high-sugar; HFHS: high-fat high-sugar). The liquids were matched in flavor (peach or blackcurrant), temperature, protein, salt and other ingredients (see^22^ for detailed liquid compositions). We used commercial skimmed milk and whole milk (British skimmed milk and British whole milk, Sainsbury’s Supermarkets Ltd., UK) as baseline low-fat and high-fat liquids and flavored the liquids with fruit juice to increase palatability

#### Nutrient foraging task

The four nutrient reward types were associated with four untrained visual cues, respectively, in each session. When a choice trial started, the monkeys were first presented with two of the four visual cues, made a touch-monitor choice between the two cues, and then received either a large amount (‘rewarded’) or a small amount (‘non-rewarded’) of the cue-associated liquids depending on its prespecified reward probability (p) (**Fig. 1A**). When the session started, two of the rewards (LFLS/HFHS or LFHS/HFLS) were offered in high reward probabilities (p=0.8), and the other two rewards in low reward probabilities (p=0.2) (**Fig. 1C**, block A or block B). The reward probabilities were reversed every 100 trials (p=0.2 **→** 0.8; p=0.8 **→** 0.2) (**Fig. 1D**).

### Data Analysis

All data were analyzed using Matlab 2017 (Mathworks).

#### Learning curve

The learning curves were plotted by aligning reward-specific choices to the probability reversal trials. In particular, based on the probability before and after reversals, we grouped these curves into incremental (P=0.2 **→** P=0.8) and decremental (P=0.8 **→** P=0.2, not shown) learning curves, and plotted the incremental curves in **Fig. 2B.**

#### Learning latency

The learning latency was defined as the number of trials between the first behavioral change point after probability reversals. The behavioral change points were identified as the significant changing points of cumulative choice slopes^49^, based on two-sample t-test with criteria P < 0.05.

#### Probability-matching (PM) choices

We simulated probability matching choices by first computing the relative proportions of the reward probabilities and transform them into predicted choices as follows^50^:

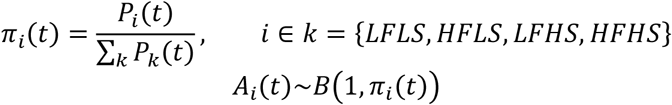

where *π_i_*(*t*) was the probability of choosing a specific option; *P_i_*(*t*) denoted the reward probability of reward *i* on trial *t*, which were summed over the stimulus set as ∑_*k*_ *P_k_*(*t*). The reward choices *A_i_*(*t*) followed the binomial distribution, based on the computed probability proportions for each reward type.

### Logistic regression analysis

#### History model

We used multiple logistic regression (*fitglm* function, Matlab) to model choices based on recent choices and reward outcomes as follows,

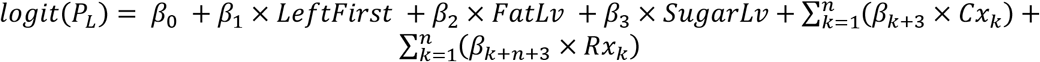

where the probability of choosing the left option (*P_L_*) was modelled by differential choice history (*Cx_n_*) and reward history (***R****x_n_*) up to recent *n* trials while controlling the presentation sequence (*LeftFirst*= 1, if the left option was shown first; 0, if the right option was shown first) and the nutrient information cued by pretrained visual stimuli (*FatLv, SugarLv* = differential fat or sugar levels = 1, if left > right; 0, if left = right; −1, if left < right). Specifically, the choice history regressors *Cx_n_* and reward history regressors *Rx_n_* were defined as the differences between the history variables of the left and right options,

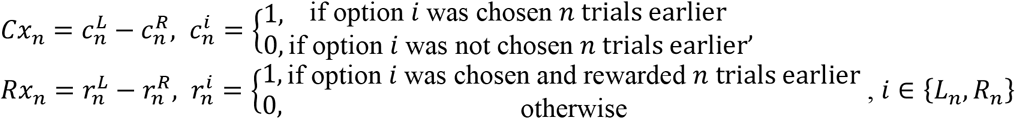

Notably, the history regressors for each option coded past trials in terms of the offered trials because the unoffered options did not carry information to influence current choices^51^. Therefore, the n-back trials for the left option may not be the same choice trials as those for the right option, due to the randomized offers.

#### Nutrient model

Based on the history model, we further included nutrient-history interaction terms to characterize the influences of fat and sugar levels on the effects of recent choices and reward outcomes:

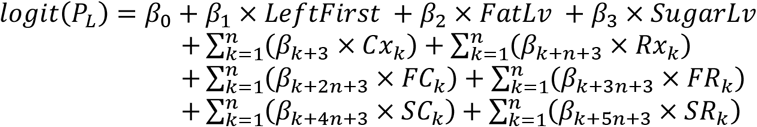

where *FC_n_* denoted recent high-fat choices and *FR_n_* for high-fat rewarded trials; *SC_n_* denoted recent high-sugar choices and *SR_n_* for high-sugar rewarded trials. The nutrient-history interaction terms were defined as follows,

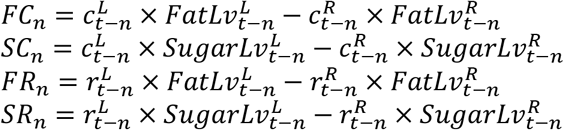

where 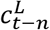 and 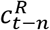 denoted whether the left or right option was chosen *n* trials earlier (1, chosen; 0, unchosen); 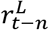 and 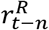 denoted whether the left or right option was chosen and was rewarded (1, chosen and rewarded; 0, otherwise).

### Reinforcement learning (RL) models

#### Standard RL model (Q-learning)

We adopted a standard Q-learning algorithm that followed the Rescorla-Wagner learning rule^1,52^. The reward values 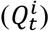 were set to be 0 for all options initially 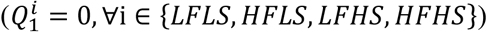 and were updated by the reward prediction errors (*RPE_t_*) multiplied by the learning rate *α* ∈ [0,1] as follows,

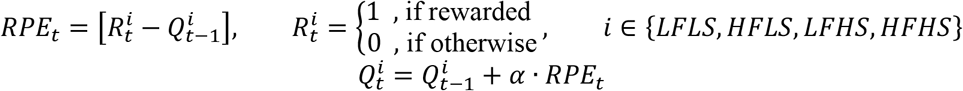

Choices were derived from transforming the value difference *δ_t_* via the softmax function into choice probability 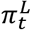, which was then dichotomized at 0.5 into binary choice actions 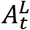 as below,

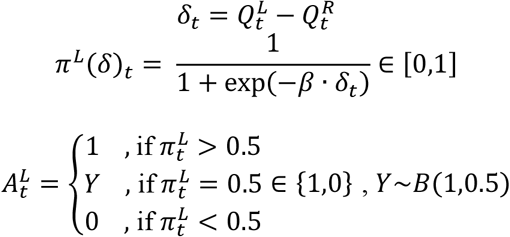

where 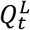 and 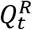 were the reward values for the left and right option on trial *t*; *β* was the inverse temperature, the sensitivity of choice to value differences.

#### Alternative RL models

We systematically included differential learning rates and nutrient-specific learning parameters into the RL models. Specifically, we examined 9 combinatorial RL models with 3 differential learning rates (*Standard*, *Asym*, and *Forget*) and 3 nutrient-specific learning parameters (*Standard*, *NutVal*, *Alpha*) (3 x 3 = 9 models) as below.

##### 1. Differential learning rates (Standard, Asym, Forget)

We included differential learning rates for rewarded (*α*^+^), unrewarded (*α*^−^), and unoffered (*α*^0^) options to update the reward values as follows,

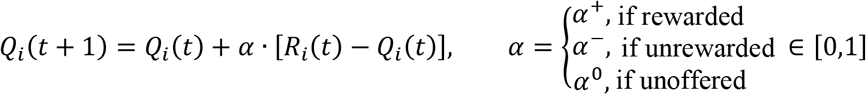

In the *Standard* model, the agent equally updated both the rewarded and unrewarded option and kept perfect memory for the unoffered option (*α*^+^ = *α*^−^, *α*^0^ = 0). In the *Asym* model, the agent updated the rewarded and unrewarded with different learning rates, while keeping perfect memory for the unoffered rewards (*α*^+^ ≠ *α*^−^, *α*^0^ = 0). In contrast, in the *Forget* model, the value of the unoffered rewards decayed due to value forgetting, but the rewarded and unrewarded option were updated equally (*α*^+^ = *α*^−^, *α*^0^ > 0).

##### 2. Nutrient-specific learning models (NutVal, Alpha)

We examined nutrient preferences by including nutient-specific values (*NutVal*) or nutrient-specific learning rates (*Alpha*). In the *NutVal* model, the reward values depend on the reward types as follows,

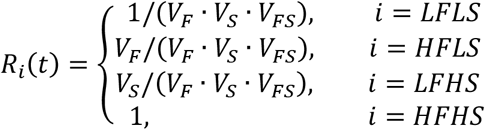

where *V_F_*, *V_S_*, and *V_FS_* are the values of high-fat content, high-sugar content, and their combinations, respectively, relative to the low-nutrient liquid. We normalized all reward values to (*V_F_* · *V_S_* · *V_FS_*), to constrain all reward values between 0 and 1.

In the *Alpha* model, higher learning rates are used to update the values for high-nutrient rewards as follow,

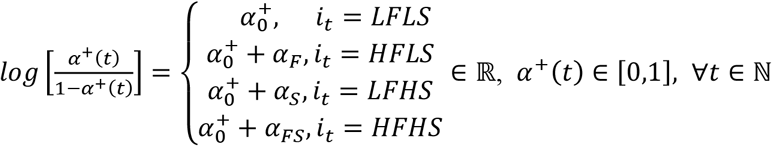

where *α*^+^(*t*) denoted the learning rate to update the value of the rewarded option on trial *t*, which was first transformed from [0,1] to any real number and modified by the high-fat level (*α_F_*), the high-sugar level (*α_S_*), or their combination (*α_FS_*). The logistic transformation ensured that the learning rates are always between 0 and 1.

#### Nutrient prediction error-RL model (NPE-RL model)

In the NPE-RL model, we decomposed the nutrient-specific values *Q_i_*(*t*) into components of fat value 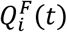 and sugar value 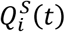,

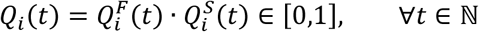

Importantly, the nutrient prediction errors were computed as the discrepancies between the subjective nutrient values and the trial-by-trial estimations of the nutrient values as follows,

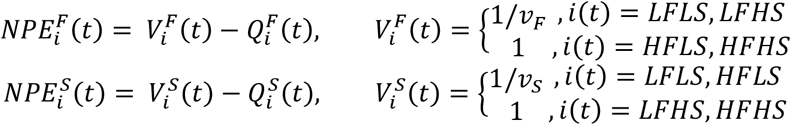

where 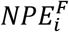 and 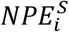 denoted the fat and sugar prediction errors for the chosen reward on trial *t*, 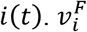 and 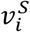 were the subjective values for fat and sugar, and 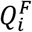 and 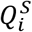 were the current values of fat and sugar components for reward *i*, respectively. The nutrient values were independently updated by corresponding nutrient prediction errors,

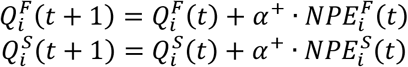

where 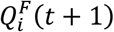 and 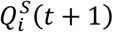 are the updated fat and sugar values, each was updated by the previous fat and sugar values, 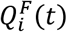 and 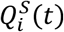, by the NPEs for fat and sugar discounted by the learning rate *α*^+^ ∈ [0,1].

## Acknowledgements

We thank Wolfram Schultz and his group for support; Putu Khorisantono for discussions; Christina Thompson and Aled David for animal care; Polly Taylor for anesthesia; Henri Bertrand for veterinary care. This work was funded by the Wellcome Trust and the Royal Society (Sir Henry Dale Fellowship 206207/Z/17/Z to F.G.). F.-Y.H. was supported by a Fellowship from the Taiwan Ministry of Education.

## SUPPLEMENTARY FIGURES

**Fig. S1.**
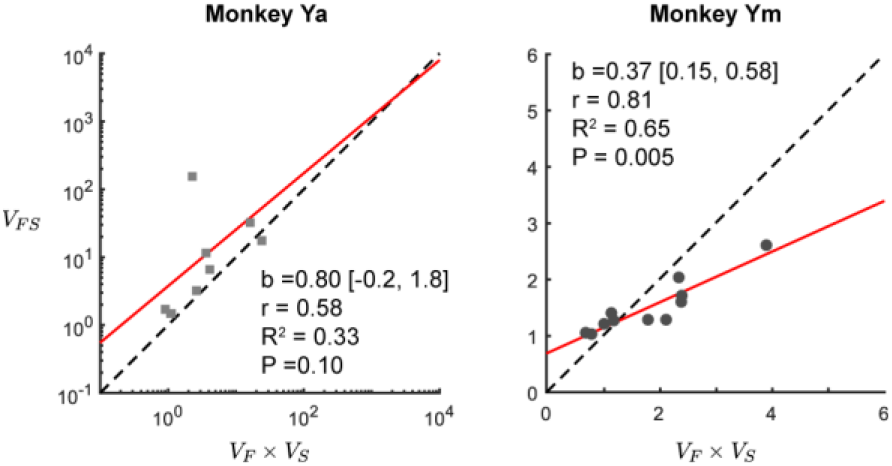
Fat-sugar value interactions. The reward values of HFHS (*V_FS_*) were plotted against the values predicted by the multiplications of the fat values (*V_F_*) and the sugar values (*V_S_*) across sessions. The unity lines (dashed) indicated the independence of the fat values and the sugar values, estimated by the nutrient-sensitive reinforcement learning models. b = slope [95% confidence interval].

**Fig. S2.**
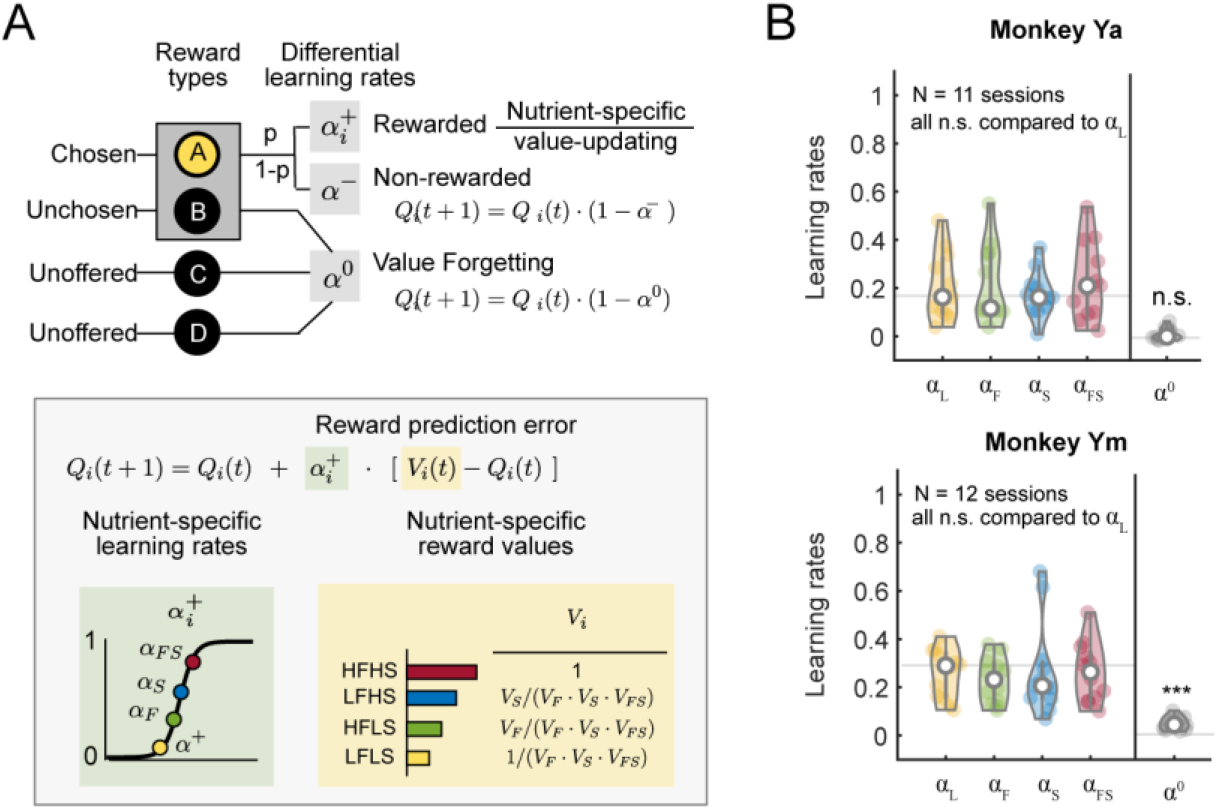
Nutrient-specific learning rates. A) Nutrient-specific learning rate model (NutAlphaVal-Forget model) architecture. The reward values were updated based on nutrient-specific learning rates 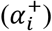 in addition to the nutrient-specific values 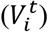. Values of the unchosen and unoffered rewards decayed according to the forgetting factor *α*^0^, as in the main nutrient value RL model (**Figure 4A**). B) Nutrient-specific learning rates and forgetting factors. Learning rates for HFLS (*α_F_*), LFHS (*α_S_*), and HFHS (*α_FS_*) were all compared to the baseline learning rates for LFLS (*α_L_*); the forgetting factors were tested against perfect value memory (*α*^0^ = 0). Wilcoxon signed-rank test.

## Notes

### Competing Interest Statement

The authors have declared no competing interest.

